# First record of a mermithid nematode (Nematoda: Mermithidae) parasitizing winged females of gall-forming aphids (Hemiptera: Aphididae: Eriosomatinae)

**DOI:** 10.1101/2021.04.10.439276

**Authors:** Xin Tong, Natsumi Kanzaki, Shin-ichi Akimoto

**Author notes:** Correspondence*: Xin Tong, Department of Ecology and Systematics, Graduate School of Agriculture, Hokkaido University, Sapporo, 060-8589 Japan.

## Abstract

Juvenile mermithid nematodes were found to parasitize winged females (sexuparae) of *Erisoma auratum* and *Tetraneura radicicola*. The morphological characteristics of mermithid nematodes are briefly described. The 18S rDNA and 28S rDNA extracted from one nematode were sequenced and used to construct a Bayesian phylogenetic tree, on which the host ranges of mermithid nematodes were represented. Our study indicated that mermithid parasitism of sexuparae led to fewer and smaller sexual female embryos. This is the first record of a mermithid in relation to eriosomatine aphids and the fourth record with respect to Aphididae.

---

Mermithid nematodes (Nematoda: Mermithidae) are obligate parasites that have been found in many invertebrates (Poinar 1975; Yeates & Buckley 2009; Kubo *et al*. 2016; Watanabe *et al*. 2021). As with other parasitic nematodes, free-living mermithid nematodes parasitize the hosts by actively penetrating the cuticles, either through natural openings of the host body, or through ingestion of their eggs by host insects (Hajek 2004). During our biological survey of eriosomatine aphids, a species of unidentified mermithid nematode was found in the abdomens of aphids collected in Hokkaido, Japan.

Aphids of Eriosomatinae (Insecta: Hemiptera: Aphididae) induce leaf galls on the primary host plants and parthenogenetically produce second-generation aphids within the gall from early May to mid-June in Hokkaido, Japan, a cool temperate zone. Second-generation aphids develop into winged adults, which migrate to the roots of secondary host plants to form colonies. In autumn, winged females (sexuparae) appear on the roots and migrate back to the primary host plants to produce sexual offspring. These offspring (male and female embryos) develop inside the abdomens of the females at their nymphal stage and are viviparously born on the trunk of the primary host plant. Sexual offspring experience both underground and aboveground environments along with their mothers from the embryonic stage until they are delivered.

After colonizing the roots of the secondary host plants, eriosomatine aphids live in the soil environment from early summer to autumn, making them susceptible to infection by soil-living parasites, such as nematodes and microbes. In the present study, we examined the rate of parasitism of the unidentified mermithid and attempted to molecularly characterize the species by employing the sequences of two ribosomal RNA genes, 18S and 28S. Thereafter, the phylogenetic status of the species in the available mermithid sequences was inferred based on the rDNA sequences.

On October 9, 2017, in Yoichi, Hokkaido, Japan (43°12′9″ N, 140°45′52″ E), autumnal winged females (sexuparae) were collected using forceps just after their alighting on the branches of *Ulmus davidiana* and maintained in 80% ethanol. Sexuparae were dissected and slide-mounted with their embryonic sexual offspring in Hoyer’s mountant for morphological observation (Tong & Akimoto 2019). When a parasite was found inside the sexuparae, it was isolated for later morphological and molecular identification. Aphids were identified morphologically, and all specimens were deposited in the Laboratory of Systematic Entomology, Hokkaido University, Sapporo, Japan.

The wing lengths of sexuparae and the body area of sexual offspring (female and male embryos) after mountant were measured and used as an index of body size (Tong & Akimoto 2019). All images were captured using a microscope eyepiece camera (Dino-Eye, AnMo Electronic Corporation, Taipei) and measurement was carried out using IMAGEJ software (http://rsbweb.nih.gov/ij/). Statistical analysis was performed using JMP software ver. Pro 14.

The isolated nematodes in mounted specimens with the host aphids were observed using light microscopy (Eclipse 80i, Nikon, Tokyo) with DIC optics and photographed with a digital camera system (MC170 HD, Leica, Wetzlar) attached to the microscope. The digital photographs were edited to enhance brightness and contrast in order to construct a micrographic figure (Fig. 1) using PhotoShop 2019 (Adobe).

**Figure 1.**
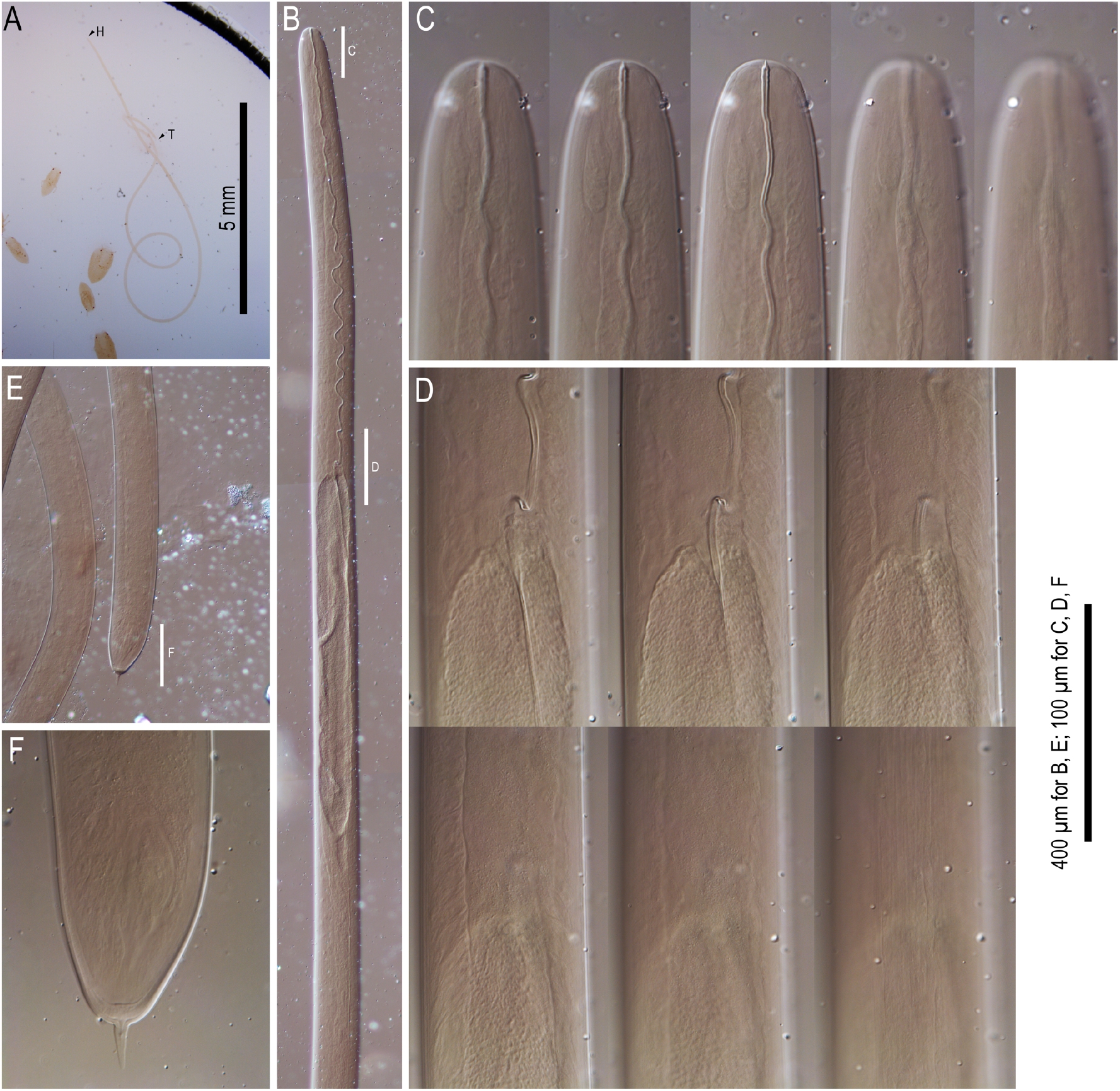
Typological characters of nematode isolated from *T. radicicola*. **A**: Whole body; **B**: Anterior region; **C**: Close-up of anterior end (“C” in subfigure B) in five different focal planes showing stoma and glands; **D**: Close-up of pharynx-intestional junction region (“D” in subfigure B) in six different focal planes showing funnel-shaped cardia and body surface structure; **E**: Posterior end of body; **F**: Close-up of tail tip (“F” in subfigure F) showing tail spike (appendage).

One parasite found in an *Eriosoma auratum* sexupara was isolated, and its genomic DNA was extracted and purified using the DNeasy Blood and Tissue Kit (QIAGEN, Venlo, the Netherlands). The 18S ribosomal gene and the gene fragment of the large ribosomal subunit (LSU) 28S rDNA sequence were amplified and polymerase chain reaction (PCR) was performed according to Kobylinski *et al*. (2012) and Shih *et al*. (2019). The following primers were used: 18S, 18S-F: 5′-CAAGGAC GAAAGTTAGAGGTTC-3′ and 18S-R: 5′-GG AAACCTTGTTACGACTTTTA-3′, and for 28S, LSU-F: 50– ACAAGTACCGTGAGGGAAAGTTG–30 and LSU-R: 50– TCGGAAGGAACCAGCTACTA–30 (Shih *et al*. 2019). The resulting templates were purified using a QIAquick PCR purification kit (QIAGEN Inc.) and sequenced in both directions using an ABI 3730xl Analyzer (Applied Biosystems). The resulting sequences were deposited in GenBank, and the BLASTn algorithm (Altschul *et al*. 1990) was applied to confirm the identity of the sequences.

The dataset of partial sequences of the nuclear 18S rDNA of mermithid nematodes in GenBank was searched and aligned using the MEGA X software package (Kumar *et al*. 2018). Host species were referenced to related publications and GenBank after obtaining 18S rDNA sequences of the parasitic mermithid nematodes (Table S1). Phylogenetic trees were constructed using Bayesian inference (BI) (Larget & Simon 1999) and maximum likelihood (ML) (Felsenstein 1981). The best-fit evolutionary model K2 + G + I was adopted by Mega X and used for all model-based methods (BI and ML). The Bayesian tree was constructed by MrBayes 3.2.7 (Ronquist *et al*. 2012) using a Markov chain Monte Carlo (MCMC) approach with 2 million generations, with tree sampling every 500 generations. The 1000 replicates were run for maximum likelihood (ML) bootstrap sampling using Mega X.

In total, 418 eriosomatine sexuparae, consisting of eight species of two genera, *Tetraneura* and *Eriosoma*, were available for examination of parasitism. Five sexuparae of *E. auratum* and one of *T. radicicola* out of the 418 individuals were found to be parasitized by a slender worm (Table 1). One parasite coexisted with embryonic sexual offspring inside the abdomen of each parasitized aphid. One of the parasites was isolated for molecular identification. The others were individually maintained with the host sexuparae in the mounted specimens for morphological observation.

**Table 1.**
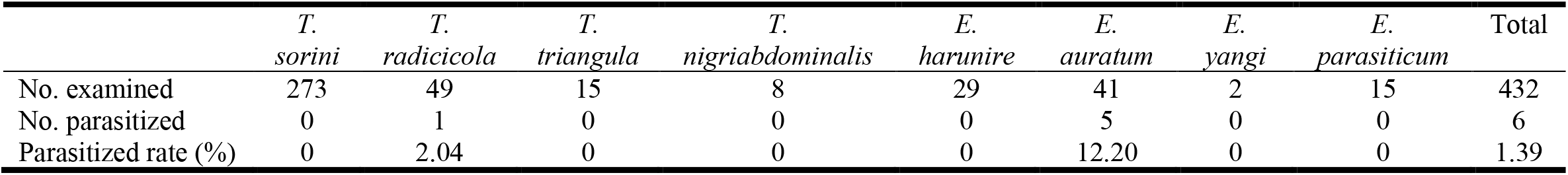
Proportion of mermithid parasitism in eriosomatine aphids collected in 2017

No significant difference was found in body size between adult mermithid-parasitized and uninfected *E. auratum* sexuparae (ANOVA, df = 1,32, *F* = 0.935, *P* = 0.34). However, the number and body size of sexual female embryos were significantly reduced in mermithid-parasitized sexuparae (df = 1,32, *F* = 9.93, *P* = 0.0035; and df = 1,32, *F* = 16.87, *P* = 0.0003, respectively) compared to uninfected sexuparae, whereas no such significant associations were found in male embryos (df = 1,32, *F* = 0.15, *P* = 0.70; and df = 1,32, *F* = 0.26, *P* = 0.61, respectively).

All nematodes were post-parasitic juveniles. One specimen that emerged from a *T. radicicola* sexupara was in relatively good condition and was examined under a stereo microscope for typological characters (Figure 1).

All isolated nematodes were juveniles without generic or species-specific characters, and some parts of the morphological structures were vague, likely because of the Hoyer fixation. Some morphological characteristics were confirmed in the specimens. The body was slender, approximately 1.5 cm long, with a smooth surface. Anterior end dome-shaped cephalic or labial papillae were not observed, possibly because of material conditions. Stoma was conspicuous, and a stylet-like well-sclerotized stoma reached the anterior end; the pharyngeal tube possessed a conspicuous lumen, connecting the stoma and cardia, and at least two gland-like structures were observed on both sides of the stoma and the anterior part of the pharyngeal tube. The cardia was funnel-shaped. Genital anlage was not confirmed, possibly because of the material conditions. The posterior end of the intestine was inconspicuous, and the anus and rectum were not observed, also likely due to the material condition. A short and bluntly pointed spike-like projection was observed at the tail tip.

The 18S rDNA and 28S rDNA gene fragments of the isolated parasite were successfully sequenced from one individual, and the sequences were deposited in GenBank under accession numbers MW649131 and MW653323. After alignment, 18S rDNA of 42 taxa and 563 base pairs were available for phylogenetic analysis. The BLAST search in GenBank indicated that the amplified sequence had the closest match and formed a clade with a previously sequenced mermithid juvenile 18S sequence (AY919185), which was collected from a grassland soil sample from Lincoln, Nebraska (Posers, pers. comm.).

Although GenBank reference sequences are limited for mermithid nematodes, here, the Bayesian-based phylogeny was constructed using currently available 18S rDNA sequences with information on the host range. Mermithid nematodes have broad host ranges, including 12 invertebrate genera, mainly Diptera and Hemiptera (Fig. 2). The mermithid sp., which was isolated from an aphid (Insecta: Hemiptera) in the present study, formed a clade with an environmental sample and was clearly separated from neighboring hemipteran associates (Fig. 2).

**Figure 2.**
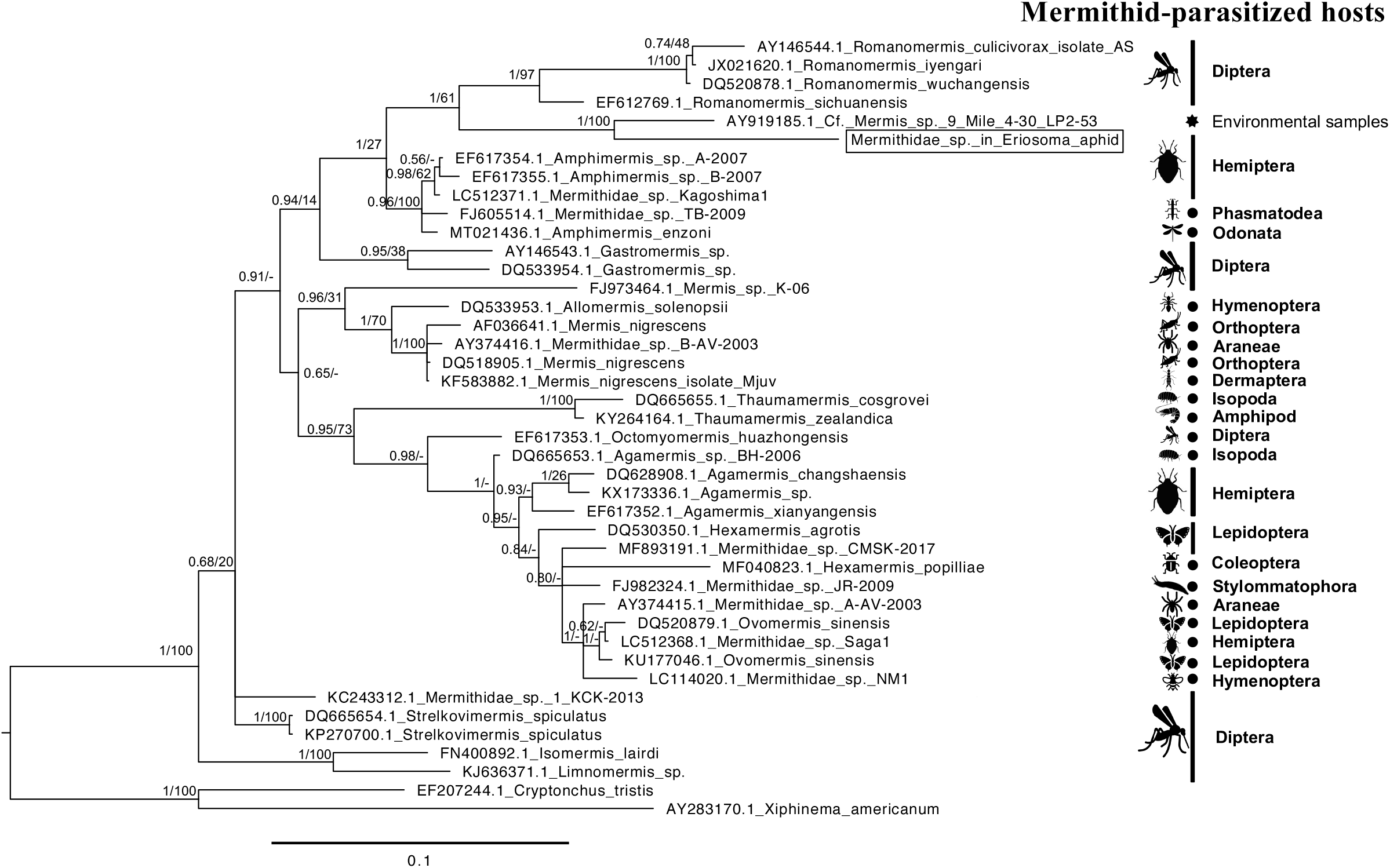
Bayesian phylogenetic tree inferred from the 18S rDNA sequences of mermithid nematodes. Values on nodes represent posterior probabilities for Bayesian inference and bootstrap support for maximum likelihood, respectively. The orders of the hosts parasitized by mermithid nematodes are listed on the right of the tree in accordance with the record of parasites.

For aphids and other herbivorous hemipteran insects that share a common arrangement of sucking mouthparts, mermithid nematodes cannot enter host bodies through mouthparts. In the present study, the unidentified mermithid nematode likely parasitized the aphid by penetrating the cuticle or gaining entry through a natural opening such as the anus. Root aphids are sedentary and susceptible to infection by nematodes and other pathogens.

Mermithid parasitism of aphids is not commonly known and only three cases have been reported (Guercio 1899; Davis 1916; Poinar 2017), although this could be due to undersampling of the aphids for this condition. The most remarkable record is the parasitism of an extinct aphid, *Caulinus burmitis* (Hemiptera: Burmitaphididae) by a fossil mermithid, which was found in mid-Cretaceous Myanmar amber (Poinar 2017). This example implies that the parasitic association between aphids and mermithid nematodes has continued for more than 100-million years. In Italy, nymphs and winged adults of the root aphid *Trama radices* Kaltenbach were found to be parasitized by an unidentified mermithid in April and May 1899, which was dispersed and embedded in the winged aphid (Guercio 1899). Davis (1916) conducted fieldwork to collect mermithid-parasitized aphids in Indiana, USA between mid-September and October 1911, and found mermithid-parasitized apterous viviparous and oviparous aphids of an *Anoecia* sp. on October 16th and 19th on the roots of *Muhlenbergia*. This is also the first record of mermithid parasitism in oviparous aphids.

The unidentified mermithid found in the present study was closest to a species collected from grassland soil around the root system of Leadplant, *Amorpha canescens* Pursh in the USA (Powers, pers. comm., also described in https://nematode.unl.edu/mermissp.htm), which possibly contained herbivorous insects, including aphids. However, the taxonomic status of the nematodes is unknown in both cases since the samples were juveniles not closely aligned to any identified species. In addition, although these two species formed a well-supported clade in the phylogenetic analysis (Fig. 2), they were clearly separated from each other considering the branch length between them. Therefore, they possibly represent separate, undersampled clades. Further collections followed by phylogenetic analyses are required to understand their relationships and taxonomic status.

The survival and performance of parasites can be largely affected by their hosts. Nematodes receive nutrition from the host tissues and hemolymph, competing with the host for nutrients that are important for its physiological development and reproduction (Smith *et al*. 1985; Mcrae *et al*. 2015). Once mermithid nematodes parasitize host insects, they can manipulate host behavior for their own benefits. For example, Allahverdipour *et al*. (2019) reported that mermithid-parasitized female mosquitoes seek water three times more than a blood source, whereas uninfected females were twice as likely to seek blood than water. Moreover, parasitizing adult hosts could be a dispersal strategy for mermithid nematodes (Campos & Sy 2003; Di Battista *et al*. 2015). In the present study, obvious morphological or behavioral alterations were not confirmed in parasitized aphids and parasitism was not detected until dissection. Nevertheless, our study indicated that mermithid parasitism in sexuparae led to fewer and smaller female sexual embryos. It is not clear whether the parasites negatively affect offspring fitness by competing for nutritious resources directly or whether maternal investment changes in response to parasitism. Thus, it is necessary to increase the sample size to investigate host manipulation by mermithid nematodes in future studies.

Mermithid nematodes can infect a broad range of aquatic and terrestrial invertebrates. However, because nematodes are often collected as juveniles, their identification and host specificity are difficult to evaluate. *Mermis nigrescens*, a parasite of grasshoppers, is reported to be found in other insect orders, such as Dermaptera, Coleoptera, and Lepidoptera (Poinar 1979). However, because of the difficulty in morphological identification, information on the host range needs to be confirmed by molecular barcoding analyses. In the present study, although the species status is still unknown, the molecular sequences can be regarded as a species-specific barcode for taxonomic identification and evaluation of the host range in future studies.

## ACKNOWLEDGEMENTS

We thank Dr. Ryoji Shinya, Meiji University, for suggestions on designing the experiments and Dr. Yuuki Kobayashi, National Institute for Basic Biology, for suggestions on phylogenetic analyses. This research was supported partially by Grant-in-Aid (19K06848) for Scientific Research from the Japan Society for the Promotion of Science (to SA) and partially by a research grant from The Yanmar Environmental Sustainability Support Association (to XT). XT is grateful to The Asahi Glass Foundation for PhD scholarship.

## SUPPORTING INFORMATION

Additional Supporting Information may be found online in the Supporting Information section at the end of the article.

**Table S1**. List of GenBank accession numbers, sample species, and references for host information included in the phylogenetic tree. DS: direct submission to GenBank.

## Notes

### Competing Interest Statement

The authors have declared no competing interest.

