## Supplementary material for "First record of a mermithid nematode (Nematoda: Mermithidae) parasitizing winged females of gall-forming aphids (Hemiptera: Aphididae: Eriosomatinae)": Table S1.: Supplementary Files.pdf

| GenBank accession no. | Sample species | References for host information |
|---|---|---|
| AY146544.1 | *Romanomermis culicivorax* | Shamseldean & Platzer 1989 |
| JX021620.1 | *Romanomermis iyengari* | Suman *et al.* 2012 |
| DQ520878.1 | *Romanomermis wuchangensis* | Duan *et al.* 2016 |
| EF612769.1 | *Romanomermis sichuanensis* | Peng *et al.* 2002 |
| AY919185.1 | *Mermithidae sp.* | DS |
| EF617354.1 | *Amphimermis sp.* | Xu *et al.* 2005 |
| EF617355.1 | *Amphimermis sp.* | Xu *et al.* 2005 |
| LC512371.1 | *Mermithidae sp.* | Iryu *et al.* 2020 |
| FJ605514.1 | *Mermithidae sp.* | Yeates & Buckley 2009 |
| MT021436.1 | *Amphimermis enzoni* | Rusconi *et al.* 2020 |
| AY146543.1 | *Gastromermis sp.* | Nickle 1972 |
| DQ533954.1 | *Gastromermis sp.* | Poinar *et al.* 2007 |
| FJ973464.1 | *Mermis sp.* | DS |
| DQ533953.1 | *Allomermis solenopsii* | Poinar *et al.* 2007 |
| AF036641.1 | *Mermis nigrescens* | Gordon & Webster 1971 |
| AY374416.1 | *Mermithidae sp.* | Vandergast & Roderick 2003 |
| DQ518905.1 | *Mermis nigrescens* | Gordon & Webster 1971 |
| KF583882.1 | *Mermis nigrescens* | Presswell *et al.* 2015 |
| DQ665655 | *Thaumamermis cosgrovei* | Tang & Hyman 2007 |
| KY264164.1 | *Thaumamermis zealandica* | Poinar *et al.* 2002 |
| EF617353.1 | *Octomyomermis huazhongensis* | Wang *et al.* 2007 |
| DQ665653.1 | *Agamermis sp.* | DS |
| DQ628908.1 | *Agamermis changshaensis* | Xu *et al.* 2005 |
| KX173336.1 | *Agamermis sp.* | Stubbins *et al.* 2016 |
| EF617352.1 | *Agamermis xianyangensis* | Kubo *et al.* 2016 |
| DQ530350.1 | *Hexamermis agrotis* | Li *et al.* 1993 |
| MF893191.1 | *Mermithidae sp.* | Kumar *et al.* 2018 |
| MF040823.1 | *Hexamermis popilliae* | Mazza *et al.* 2017 |
| FJ982324.1 | *Mermithidae sp.* | Ross *et al.* 2010 |
| AY374415.1 | *Mermithidae sp.* | Vandergast & Roderick 2003 |
| DQ520879.1 | *Ovomermis sinensis* | Sun *et al.* 2020 |
| LC512368.1 | *Mermithidae sp.* | Iryu *et al.* 2020 |
| KU177046.1 | *Ovomermis sinensis* | Sun *et al*, 2020 |
| LC14020.1 | *Mermithidae sp.* | Kubo *et al.* 2016 |
| KC243312.1 | *Mermithidae sp.* | Kobylinski *et al.* 2012 |
| DQ665654.1 | *Strelkovimermis spiculatus* | Poinar & Camino 1986 |
| KP270700.1 | *Strelkovimermis spiculatus* | Poinar & Camino 1986 |
| FN400892.1 | *Isomermis lairdi* | Gradinarov D 2014 |
| KJ636371.1 | *Limnomermis sp.* | Villemant *et al.* 2015 |
| EF207244.1 | *Cryptonchus tristis* | Holterman *et al.* 2008 |
| AY283170.1 | *Xiphinema americanum* | Neilson *et al.* 2004 |
